# Do water and host size interactively affect the impact of a native hemiparasite on a major invasive legume?

**DOI:** 10.1101/2022.01.22.477374

**Authors:** Robert M. Cirocco, Evelina Facelli, José M. Facelli

## Abstract

It is unknown how the combined effects of host size and water availability influence parasitic plant:host associations. This is a major gap considering that parasitic plants would frequently encounter hosts of different size growing in different water conditions in nature. In a glasshouse experiment, small or large *Ulex europaeus* (major invasive host) were provided high or low water supply and infected or not with the Australian native shoot hemiparasitic vine, *Cassytha pubescens*. Infection significantly decreased host total, shoot and root biomass, in both low water and high water conditions but more severely so in the latter. Infection also significantly decreased total biomass of both large and small plants and more severely so for the latter. Infection significantly decreased host foliar nitrogen and potassium concentrations in well-watered but not in low water conditions. Infection significantly suppressed host predawn and midday quantum yield, midday electron transport rates, foliar phosphorus concentration and nodule biomass, irrespective of water conditions or host size. Parasite biomass (including g^-1^ host total biomass) was significantly greater on hosts growing in well-watered than in low water conditions. Our results suggest that some native parasitic plants may effectively control major invasive hosts, particularly in wetter habitats and or when the host is small, leading to enhanced biodiversity protection in those settings.

## Introduction

Parasitic plants are an important global group of approximately 4500 species that can have significant impacts on ecosystems, communities and plant populations (Těšitel et al., 2021). For example, the root hemiparasite *Rhinanthus minor* has a greater effect on grasses (dominant species) than forbs, resulting in a significant increase in plant diversity and abundance of total invertebrates (Hartley et al., 2015). Parasitic plants can also have greater effects on major invasive weeds than native hosts, and thus show potential as weed biocontrol agents for biodiversity protection (Těšitel et al., 2020). For instance, in China, the native holoparasitic vine *Cuscuta australis* was found to infect invasive hosts much more than native hosts, leading to decreased cover of invasive hosts and a significant increase in plant diversity (Yu et al., 2011). Also, in Australia, the native hemiparasitic vine *Cassytha pubescens* has been found to have a greater impact on major invasive weeds than on native hosts studied (Prider et al., 2009; Shen et al., 2010; Cirocco et al., 2016a, 2017, 2018). At least for the *R. minor* and *C. pubescens* examples above, the greater parasite effect on dominant or invasive hosts may be explained by the parasite’s haustoria connecting more effectively to the vasculature of these hosts (relative to sub-dominant or native hosts) which leads to increased resource removal and impact (Cameron and Seel, 2007; Facelli et al., 2020)

Upon infection, there are a number of biotic (e.g. host size) and abiotic factors (e.g. water availability) that may influence the degree of impact a parasitic plant has on a particular host. There are only two studies that have investigated how host size or age modulates the impact of parasitic plants. In China, *Cuscuta australis* was found to have a negative impact on the invasive host *Bidens pilosa*, but only when it was younger and smaller (Li et al., 2015). In Australia, *C. pubescens* was found to have a negative effect on the invasive host *Ulex europaeus*, but more severely so when the host plants were smaller (Cirocco et al., 2020). Also, the anomaly that *C. pubescens* negatively affected the native host *Acacia paradoxa* was attributed to this host being smaller than in a previous experiment where this host was unaffected by the parasite (Cirocco et al., 2017, 2021a).

As water is essential for plants and is a main resource removed by parasites, it is paramount that we understand how its availability may affect parasite:host associations, especially in Mediterranean climates that are expected to become increasingly drier as a result of climate change (Klausmeyer and Shaw, 2009). Yet, there are surprisingly few studies that have investigated the influence of water on parasite effects on their hosts (e.g. Těšitel et al., 2015; Korell et al., 2019). One study found that the negative effect of *Cuscuta australis* on stomatal conductance and transpiration of the invasive weed *Mikania micrantha* was more severe in low water conditions (Le et al., 2015). In contrast, *Cuscuta gronovii* grew better and had a stronger impact on host growth in well-watered conditions (Evans and Borowicz, 2013, 2015). Cirocco et al. (2016b) also found that *C. pubescens* was larger and had a more severe impact on performance of *U. europaeus* in high water than low water treatments. With impending climate change and the research above highlighting how both host size and water supply can greatly change the degree of parasite impact, it would be important to investigate their combined effects on parasitic plant:host associations. However, to the best of our knowledge, there are no such studies. This is a major gap in the literature considering that, in nature, parasitic plants would frequently encounter hosts of differing size in areas that vary in water availability.

Here, we investigated the combined influence of host size and water on the performance of the native parasitic vine *C. pubescens* and its impact on the major invasive weed *U. europaeus*. Based on previous findings mentioned, we hypothesised that the native parasite would have the greatest impact on smaller hosts growing in well-watered conditions. We also expected the parasite to grow best on larger hosts growing in well-watered conditions, considering we have found the parasite grows better on larger hosts (Cirocco et al., 2020) or in high water conditions (Cirocco et al., 2016b). A number of key plant traits were measured to assess the performance of the parasite and its impact on the host in the various treatments: 1) chlorophyll fluorescence parameters (i.e. predawn and midday quantum yield: *F*_v_/*F*_m_, Φ_PSII_, respectively) provide deep insights in to how efficiently the plant is utilising light for photosynthesis; 2) carbon isotope composition is a powerful long-term indicator of how conservative the plant is with its water-use relative to the amount of carbon being fixed, with less negative values representing a higher water-use efficiency (Lambers et al., 2008); 3) plant nutrient-status is important because it can help explain higher-order changes in both photosynthesis and growth of the parasite and host and 4) growth measurements (e.g. biomass, rhizobial nodulation).

## Materials and methods

### Study species

*Ulex europaeus* L. (Fabaceae) is a perennial, evergreen spiny shrub, that can reach 4 m in height, live for approximately 30 years and is found in areas with annual rainfall of around 650–900 mm (Richardson and Hill, 1998). It forms associations with N_2_-fixing bacteria, providing carbohydrate to rhizobial strains such as *Bradyrhizobium*, in exchange for reduced nitrogen (Weir et al., 2004; Rodríguez-Echeverría, 2010). *Ulex europaeus* produces large amounts of seed that remain viable in the soil for decades and quickly colonises areas after disturbance (Parsons and Cuthbertson, 2001). It originated in the Iberian Peninsula and then radiated into northern Europe, following introduction into most parts of the world it has been classified as one of the world’s 100 worst invasive species (Lowe et al., 2000; see Hornoy et al., 2013). *Cassytha pubescens* R. Br. (Lauraceae) is an Australian native perennial hemiparasitic vine which has photosynthetic stems (approximately 0.5–1.5 mm in diameter) with indeterminate growth that coil around and attach to shoots of multiple hosts with multiple ellipsoid haustoria (Weber, 1981; Kokubugata et al., 2012). It is a generalist parasite naturally infecting both native and invasive shrubs including *U. europaeus* (Cirocco pers. obs.).

### Experimental design

*Ulex europaeus* seeds were placed in near boiling water and left to soak overnight and then sowed in growing tubes containing Waikerie sand (pH ~7) in late February 2020. In late July (2020), the seedlings were transplanted into 0.8-l pots (90 mm Width x 180 mm Height) containing the same soil, and they were categorised as being small or larger (approximately 20% greater) based on their differences in height (Figure S1A). Then, in late October (2020) these small (S) and large (L) plants (*n* = 32) were randomly allocated into treatments and 8 blocks: uninfected (-) or infected (+) with *C. pubescens;* grown in well-watered (W) or low water (D) conditions, with each block containing a single replicate from all treatment combinations i.e. Block 1 = 1SW-, 1SW+, 1LW-, 1LW+, 1SD-, 1SD+, 1LD-, 1LD+. Watering treatments were not imposed (i.e. plants were well watered) until plants were successfully infected with the parasite. Plants were infected using the technique of Shen et al. (2010), where *U. europaeus* already infected with *C. pubescens* were placed adjacent experimental plants. The parasite being a vine with indeterminate growth coiled around and attached to plants randomly designated for infection. Parasite tendrils infecting one plant were prevented from coiling around and attaching to other plants in order to obtain separately infected individuals. The next step was to cut the parasite connection between the ‘donor’ plants and experimental plants, however, if cut too early, the parasite would not have haustorially connected to and established on the experimental plants, thus jeopardising the experiment. There is no definitive way of knowing if the parasite has successfully established, so the timing of cutting the connection errs on the side of caution and is based on over 10 years of experience using this technique. The infection process began in early November 2020 and was completed by late January 2021.

At this time, plant height was measured again (Figure S1B,C,D). Then, uninfected and infected small and large plants were placed into their allocated blocks and watering treatments were imposed from early February 2021 to mid-April 2021. Well-watered and low water plants were maintained at 100% and 60% field capacity respectively, by weighing plants daily and watering accordingly to weight. Field capacity of the soil and corresponding weight for watering was determined by a modified version of the filter paper technique (Bouyoucos, 1929; see Cirocco et al., 2016b for details). Low water plants were kept at 60% field capacity because at lower values the parasite would wilt (Cirocco pers. obs.). The experiment was conducted in an evaporatively cooled glasshouse at The University of Adelaide, plants were re-randomised within blocks fortnightly to negate any small light differences. Uninfected and infected plants in the different treatments were all supplied with liquid fertiliser (Nitrosol, Rural Research Ltd, Auckland, New Zealand; NPK 8:3:6) monthly at the manufacturer’s recommended dosage.

### Chlorophyll fluorescence, biomass, δ^13^C and nutrient concentration

Predawn (*F*_v_/*F*_m_) and midday quantum yield (Φ_PSII_) and midday electron transport rates (ETR) of *U. europaeus* (*n* = 6–8) and *C. pubescens* (*n* = 6–8) were measured 67 and 66 days after treatments were imposed (DAT), respectively, using a portable chlorophyll fluorometer (MINI-PAM, Walz, Effeltrich, Germany) equipped with a leaf-clip (2030–B, Walz, Effeltrich, Germany). Midday yield and ETR measurements of *U. europaeus* and *C. pubescens* were made on a sunny day between 12:00–13:40 h at a photon flux density of 1004 ± 7 μmol m^-2^ s^-1^ (*n* = 87). At the end of the experiment (Figures S2–S5), plants (including parasite) from all eight blocks were harvested, minus six plants (two plants each from LD−, SD− and SD+ treatments) that died from an unknown reason (i.e. *n* = 6–8). Due to limited oven space: above and below ground biomass (including nodules) of four blocks were harvested 69–71 DAT, and the remaining four blocks were harvested 77–78 DAT. Harvested material was oven-dried at 60°C for seven days. Carbon isotope composition (δ^13^C) and nitrogen [N] concentration of dried spines and stems of *U. europaeus* and *C. pubescens*, respectively, (*n* = 6), were quantified by mass spectrometry (GV Instruments, Manchester, UK). Potassium [K] and phosphorus [P] concentrations of dried spines of *U. europaeus* and parasite stems (*n* = 6) were determined using inductively coupled plasma spectroscopy (Cuming Smith British Petroleum Soil and Plant Laboratory, Western Australia).

### Statistical analysis

The influence of infection, water and host size on performance of *U. europaeus* was analysed with three-way ANOVA (block as random effect). The influence of water and host size on parasite performance was analysed with a two-way ANOVA (block as random effect). For any significant interactions we used pairwise comparisons within each level of the other effect(s) in the model using a Tukey HSD test. If an interaction was not found, we considered main effects of infection, water or size valid. For example, a main effect of infection compares between uninfected (SW−, LW−, SD−, LD− plants pooled) and infected plants (SW+, LW+, SD+, LD+ plants pooled) for that particular variable. Model assumptions were met, in some cases this involved data transformation where stated and all data were analysed with the package lme4 in R (R Development Core Team, 2016) and α = 0.05 (Type 1 error rate).

## Results

### Host and parasite growth

Infection and water had an interactive effect on total, shoot and root biomass of *U. europaeus*, irrespective of host size (Table 1, no three-way interactions: Figure 1A,D,F). Total, shoot and root biomass of infected plants in well-watered and low water conditions were approximately 70% and 60% lower than those of uninfected ones, respectively (Figure 1B,E,G). Infection and host size also had an interactive effect on host total biomass, regardless of watering treatments (Table 1). Total biomass of small and large infected plants were 70% and 66% lower relative to that of small and large uninfected ones, respectively (Figure 1C). Water and size had an interactive effect on host total and root biomass, irrespective of host infection-status (Table 1). Total biomass of large low water plants was 33% lower than that of large well-watered plants, whereas total biomass of small plants was unaffected by watering treatment (Figure S6A). Root biomass of large low water plants was 34% lower than that of large well-watered plants, whereas root biomass of small plants was unaffected by watering treatment (Figure S6B). There was a main effect of host size on host shoot biomass which was 21% lower on small plants relative to large ones (Table 1; Figure S6C).

**TABLE 1.** Three-way ANOVA results for the effects of infection with *Cassytha pubescens* (I), water (W) and host size (S) on total, shoot and root biomass, shoot:root ratio (S:R), nodule biomass (Nod) and nodule biomass g^-1^ root biomass (Nod g^-1^ root) of *Ulex europaeus*

|  | Total | Shoot | Root | S:R | Nod | Nod g<sup>-1</sup> root |
| --- | --- | --- | --- | --- | --- | --- |
| **I** |  |  |  |  |  |  |
| *P*-value | **<0.0001\*\*\*** | **<0.0001\*\*\*** | **<0.0001\*\*\*** | **0.005\*\*** | **<0.0001\*\*\*** | **<0.0001\*\*\*** |
| *F*-value | 540 | 541 | 195 | 8.57 | 19.4 | 33.7 |
| *df* | 1, 44.7 | 1, 44.5 | 1, 45.3 | 1, 45.3 | 1, 44.7 | 1, 44.5 |
| **W** |  |  |  |  |  |  |
| *P*-value | **<0.0001\*\*\*** | **<0.0001\*\*\*** | **0.007\*\*** | **0.986** | **0.005\*\*** | **0.268** |
| *F*-value | 28.5 | 30.5 | 8.04 | 0.0003 | 8.85 | 1.26 |
| *df* | 1, 44.0 | 1, 43.9 | 1, 44.5 | 1, 44.5 | 1, 44.0 | 1, 43.2 |
| **I × W** |  |  |  |  |  |  |
| *P*-value | **<0.0001\*\*\*** | **<0.0001\*\*\*** | **0.033\*** | **0.004\*\*** | **0.503** | **0.791** |
| *F*-value | 29.4 | 35.5 | 4.82 | 9.34 | 0.455 | 0.071 |
| *df* | 1, 45.2 | 1, 45.0 | 1, 45.9 | 1, 45.9 | 1, 45.2 | 1, 45.4 |
| **S** |  |  |  |  |  |  |
| *P*-value | **<0.0001\*\*\*** | **<0.0001\*\*\*** | **<0.0001\*\*\*** | **0.758** | **0.002\*\*** | **0.631** |
| *F*-value | 42.4 | 36.3 | 23.2 | 0.096 | 10.4 | 0.235 |
| *df* | 1, 43.5 | 1, 43.4 | 1, 43.8 | 1, 43.8 | 1, 43.5 | 1, 43.0 |
| **I × S** |  |  |  |  |  |  |
| *P*-value | **0.038\*** | **0.076** | **0.069** | **0.313** | **0.840** | **0.977** |
| *F*-value | 4.59 | 3.31 | 3.49 | 1.04 | 0.041 | 0.0008 |
| *df* | 1, 43.3 | 1, 43.3 | 1, 43.6 | 1, 43.6 | 1, 43.4 | 1, 42.6 |
| **W × S** |  |  |  |  |  |  |
| *P*-value | **0.009\*\*** | **0.056** | **0.005\*\*** | **0.146** | **0.662** | **0.166** |
| *F*-value | 7.46 | 3.85 | 8.94 | 2.19 | 0.194 | 1.99 |
| *df* | 1, 43.7 | 1, 43.6 | 1, 44.0 | 1, 44.0 | 1, 43.7 | 1, 43.4 |
| **I × W × S** |  |  |  |  |  |  |
| *P*-value | **0.243** | **0.426** | **0.168** | **0.318** | **0.470** | **0.672** |
| *F*-value | 1.40 | 0.645 | 1.97 | 1.02 | 0.531 | 0.182 |
| *df* | 1, 43.5 | 1, 43.5 | 1, 43.7 | 1, 43.7 | 1, 43.5 | 1, 42.9 |

**P**, **F** and **df** values are in **bold**, *italics* and regular type, respectively, significance codes:
<0.0001-0.001'\*\*\*', 0.001-0.01'\*\*\*', 0.01-0.05'\*. S:R log transformed and Nod g<sup>-1</sup> root square root transformed to meet model assumptions.

**FIGURE 1.**
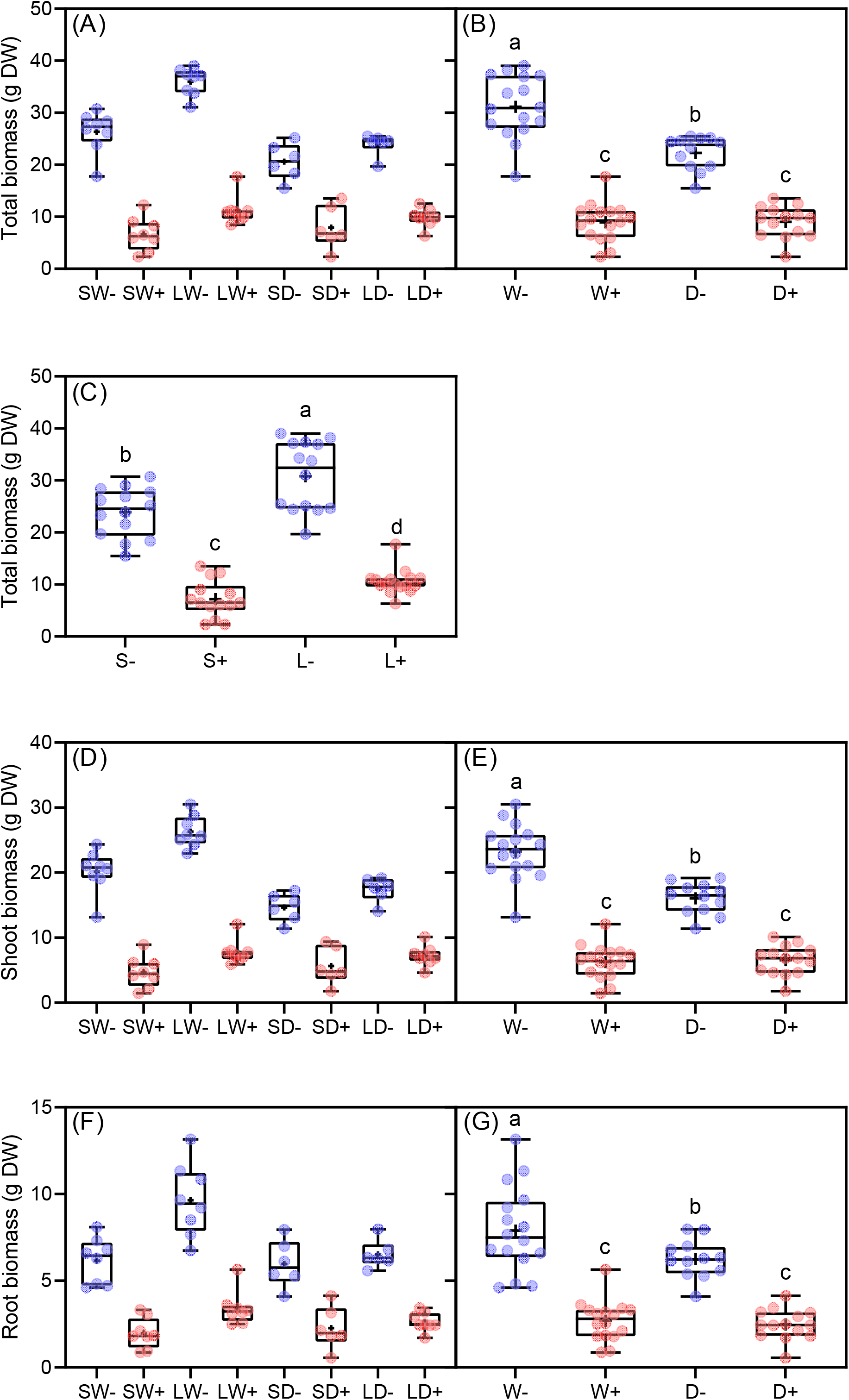
(A) Total, (D) Shoot and (F) root biomass of small (S) or large (L) *Ulex europaeus* uninfected (−) or infected (+) with *Cassytha pubescens* growing in well-watered (W) or low water (D) conditions. Infection × water interactive effect on host (B) total, (E) shoot and (G) root biomass. (C) Infection × size interactive effect on host total biomass. All data points, median, 1^st^ and 3^rd^ quartiles, interquartile range and mean (+ within box) are shown, different letters signify significant differences: (A, D, F) *n* = 8 (except SD−, LD− and SD+: *n* = 6), (B, E, G) *n* = 12–16, (C) *n* = 14–16

Infection and water had an interactive effect on the shoot:root ratio (S:R) of *U. europaeus*, regardless of host size (Table 1, no three-way interaction: Figure 2A). Shoot:root ratio of infected plants in well-watered conditions was 25% lower than that of respective uninfected plants while S:R of low water plants was unaffected by infection (Figure 2B). Main effects of infection, water and size were found for nodule biomass of *U. europaeus* (Table 1, no threeway interaction: Figure 2C). Nodule biomass of infected plants was 33% lower than that of uninfected ones (Figure 2D). Nodule biomass of hosts in low water conditions or small in size was approximately 25% lower than that of hosts in well-watered conditions or large in size, respectively (Figure 2E,F). There was a main effect of infection on nodule biomass g^-1^ root biomass (Table 1, no 3-way interaction: Figure 2G). Nodule biomass g^-1^ root biomass was 45% higher on infected than uninfected plants (Figure 2H). There were main effects of water or host size on biomass of *C. pubescens* (Table 2, no interaction: Figure 3A). Parasite biomass on hosts in low water was 42% lower than that on hosts in well-watered conditions (Figure 3B). Parasite biomass on small hosts was 21% lower relative to that on large ones (Figure 3C). There was a main effect of water on parasite biomass g^-1^ host total biomass (Table 2, no interaction: Figure 3D). Parasite biomass g^-1^ host total biomass in low water treatments was 39% lower than that in well-watered treatments (Figure 3E).

**FIGURE 2.**
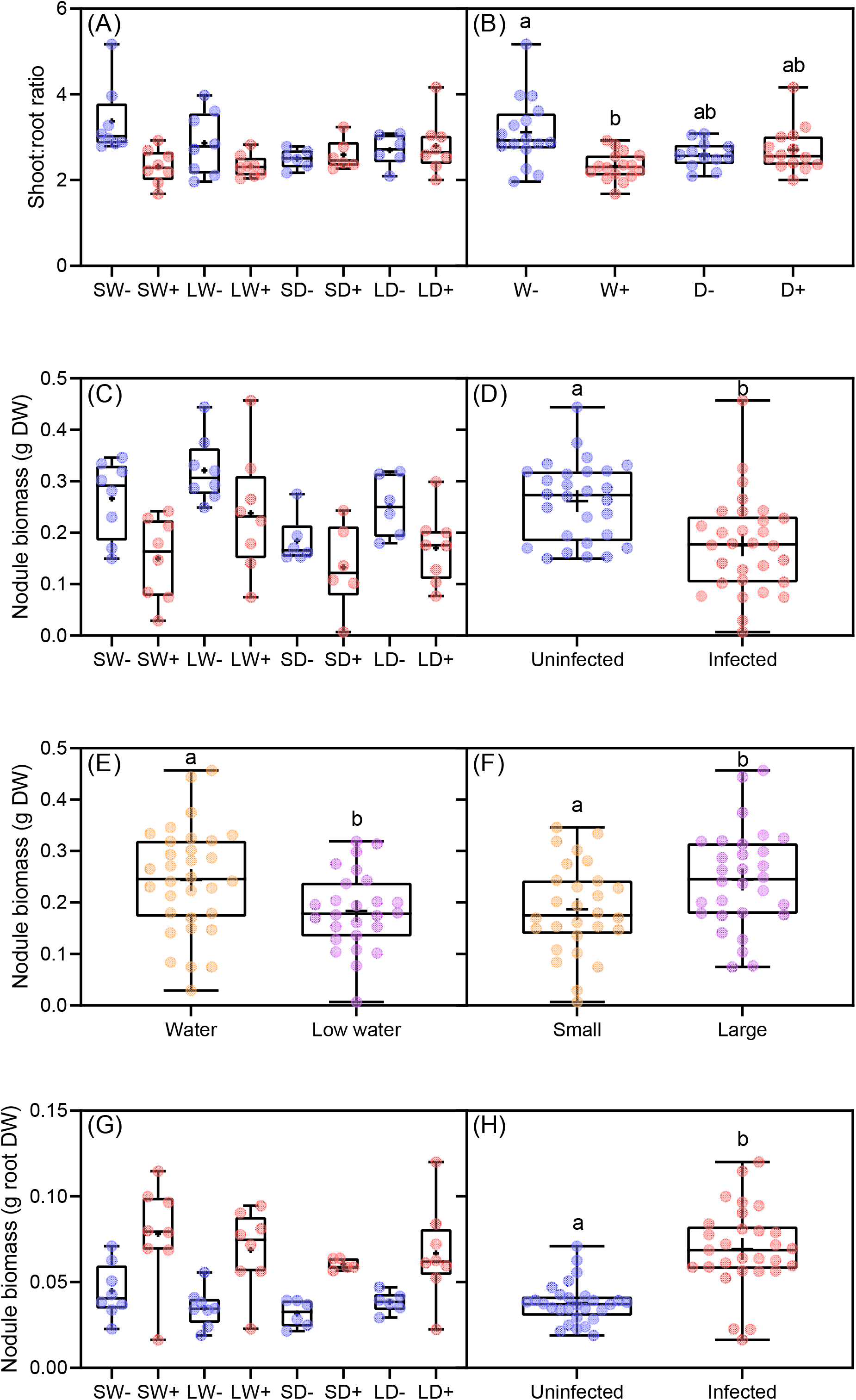
(A) Shoot:root ratio, (C) nodule biomass and (G) nodule biomass g^-1^ root biomass of small (S) or large (L) *Ulex europaeus* uninfected (-) or infected (+) with *Cassytha pubescens* growing in well-watered (W) or low water (D) conditions. (B) Infection × water interactive effect on host shoot:root ratio. Main effect of infection (D), water (E) and size (F) on nodule biomass. (H) Main effect of infection on nodule biomass g^-1^ root biomass. All data points, median, 1^st^ and 3^rd^ quartiles, interquartile range and mean (+ within box) are shown, different letters signify significant differences: (A, C, G) *n* = 8 (except SD−, LD− and SD+: *n* = 6), (B) *n* = 12–16, (D, F) *n* = 28–30, (E) *n* = 26–32 and (H) *n* = 28–29

**FIGURE 3.**
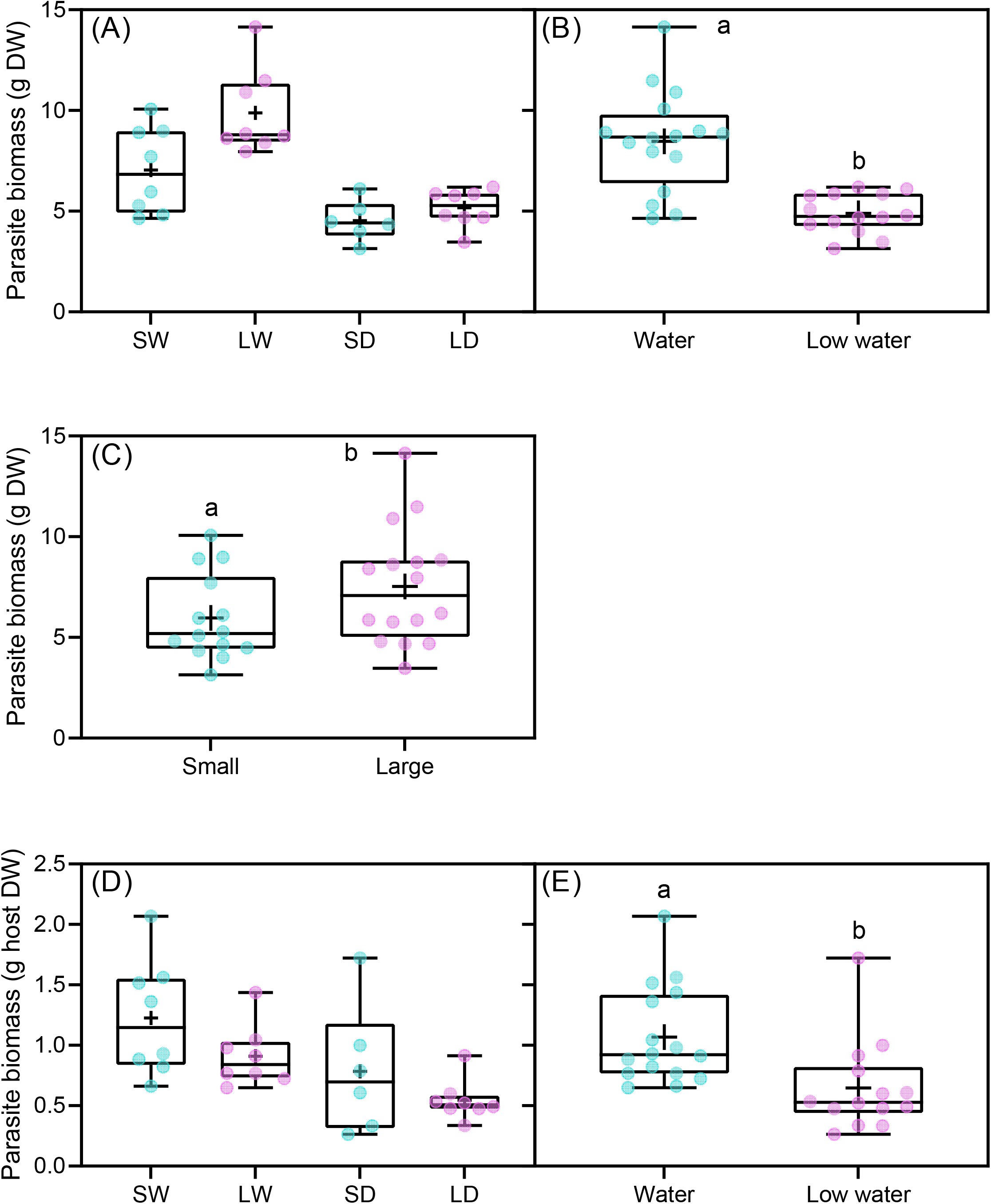
(A) Parasite biomass and (D) parasite biomass g^-1^ host total biomass of *Cassytha pubescens* when infecting small (S) or large (L) *Ulex europaeus* growing in well-watered (W) or low water (D) conditions. Main effect of (B) water and (C) size on parasite biomass and main effect of (E) water on parasite biomass g^-1^ host total biomass. All data points, median, 1^st^ and 3^rd^ quartiles, interquartile range and mean (+ within box) are shown, different letters signify significant differences: (A, D) *n* = 8 (except SD: *n* = 6) and (B, C, E) *n* = 14–16

**TABLE 2.** Two-way ANOVA results for effects of water (W) and host size (S) on parasite biomass, parasite biomass g^-1^ host total biomass, predawn and midday quantum yield (*F*_v_/*F*_m_ and Φ_PSII_), midday electron transport rates (ETR), carbon isotope composition (δ^13^C) and stem nitrogen [N], phosphorus [P] and potassium [K] concentrations of *Cassytha pubescens* when infecting *Ulex europaeus*

| | Parasite<br>biomass | Parasite<br>biomass<br>g<sup>-1</sup> host | $F_v/F_m$ | $\Phi_{PSII}$ | ETR | $\delta^{13}C$ | [N] | [P] | [K] |
| --- | --- | --- | --- | --- | --- | --- | --- | --- | --- |
| W |  |  |  |  |  |  |  |  |  |
| *P*-value | **<0.0001\*\*\*** | **0.001\*\*** | **0.124** | **0.037\*** | **0.039\*** | **<0.0001\*\*\*** | **0.708** | **0.007\*\*** | **<0.0001\*\*\*** |
| *F*-value | *47.2* | 15.0 | *2.59* | *5.01* | *4.92* | *112* | *0.147* | *10.0* | *32.7* |
| *df* | 1, 19.4 | 1, 19.5 | 1, 19.2 | 1, 19.6 | 1, 19.7 | 1, 15.4 | 1, 14.3 | 1, 14.6 | 1, 14.1 |
| S |  |  |  |  |  |  |  |  |  |
| *P*-value | **0.003\*\*** | **0.095** | **0.387** | **0.225** | **0.244** | **0.533** | **0.309** | **0.007\*\*** | **0.315** |
| *F*-value | *11.8* | 3.08 | *0.785* | *1.57* | *1.44* | *0.407* | *1.11* | *10.0* | *1.09* |
| *df* | 1, 19.4 | 1, 19.5 | 1, 19.2 | 1, 19.6 | 1, 19.7 | 1, 15.4 | 1, 14.3 | 1, 14.6 | 1, 14.1 |
| W × S |  |  |  |  |  |  |  |  |  |
| *P*-value | **0.071** | **0.891** | **0.663** | **0.689** | **0.792** | **0.677** | **0.819** | **0.357** | **0.157** |
| *F*-value | *3.65* | 0.020 | *0.196* | *0.165* | *0.071* | *0.180* | *0.054* | *0.905* | *2.24* |
| *df* | 1, 19.5 | 1, 19.5 | 1, 19.2 | 1, 19.7 | 1, 19.8 | 1, 15.5 | 1, 14.6 | 1, 14.9 | 1, 14.3 |

**P**, **F** and **df** values are in **bold**, *italics* and regular type, respectively, significance codes:
<0.0001-0.001'\*\*\*\*', 0.001-0.01'\*\*\*', 0.01-0.05'\*. Parasite biomass g<sup>-1</sup> host total biomass log transformed to meet model assumptions.

### Host and parasite photosynthetic performance, δ^13^C and nitrogen status

There was main effect of infection on host *F*_v_/*F*_m_, Φ_PSII_ and ETR of *U. europaeus* (Table 3, no interactions: Figure 4A,C,E). *F*_v_/*F*_m_, Φ_PSII_ and ETR of infected plants were 4%, 31% and 34% lower than those of uninfected ones (Figure 4B,D,F). There was also a main effect of water, with *F*_v_/*F*_m_, Φ_PSII_ and ETR of low water plants being 5%, 28% and 31% lower compared with those of well-watered plants (Table 3; Figure S7). No significant treatment effects were found for *F*_v_/*F*_m_ of *C. pubescens* (Table 2; Figure 5A). Water had a main effect on Φ_PSII_ and ETR of *C. pubescens* (Table 2, no interactions: Figure 5B,D). Φ_PSII_ and ETR of *C. pubescens* when infecting low water plants were approximately 20% lower than those when infecting well-watered hosts (Figure 5C,E).

**TABLE 3.** Three-way ANOVA results for the effects of infection with *Cassytha pubescens* (I), water (W) and host size (S) on predawn and midday quantum yield (*F*_v_/*F*_m_ and Φ_PSII_), midday electron transport rates (ETR), carbon isotope composition (δ^13^C), and foliar nitrogen [N], phosphorus [P] and potassium [K] concentrations of *Ulex europaeus*

| | $F_v/F_m$ | $\Phi_{PSII}$ | ETR | $\delta^{13}C$ | [N] | [P] | [K] |
| --- | --- | --- | --- | --- | --- | --- | --- |
| **I** |  |  |  |  |  |  |  |
| *P*-value | **<0.0002\*\*\*** | **0.0004\*\*\*** | **<0.0001\*\*\*** | **0.142** | **0.277** | **0.0007\*\*\*** | **0.011\*** |
| *F*-value | 17.2 | 14.5 | 18.3 | 2.26 | 1.22 | 13.7 | 7.28 |
| *df* | 1, 45.3 | 1, 44.9 | 1, 45.0 | 1, 35.5 | 1, 35.2 | 1, 35.5 | 1, 35.2 |
| **W** |  |  |  |  |  |  |  |
| *P*-value | **<0.0001\*\*\*** | **0.003\*\*** | **0.0009\*\*\*** | **0.0006\*\*\*** | **0.901** | **0.104** | **<0.0001\*\*\*** |
| *F*-value | 25.7 | 10.1 | 12.6 | 14.3 | 0.016 | 2.79 | 23.7 |
| *df* | 1, 44.5 | 1, 44.2 | 1, 44.3 | 1, 35.5 | 1, 35.3 | 1, 35.5 | 1, 35.3 |
| **I × W** |  |  |  |  |  |  |  |
| *P*-value | **0.620** | **0.257** | **0.348** | **0.061** | **0.006\*\*** | **0.212** | **0.036\*** |
| *F*-value | 0.249 | 1.32 | 0.899 | 3.73 | 8.53 | 1.62 | 4.77 |
| *df* | 1, 45.9 | 1, 45.5 | 1, 45.6 | 1, 35.5 | 1, 35.2 | 1, 35.5 | 1, 35.3 |
| **S** |  |  |  |  |  |  |  |
| *P*-value | **0.869** | **0.560** | **0.669** | **0.023\*** | **0.256** | **0.0004\*\*\*** | **0.402** |
| *F*-value | 0.028 | 0.345 | 0.186 | 5.62 | 1.34 | 15.6 | 0.721 |
| *df* | 1, 43.8 | 1, 43.6 | 1, 43.7 | 1, 35.5 | 1, 35.2 | 1, 35.5 | 1, 35.2 |
| **I × S** |  |  |  |  |  |  |  |
| *P*-value | **0.187** | **0.120** | **0.184** | **0.271** | **0.488** | **0.130** | **0.192** |
| *F*-value | 1.80 | 2.52 | 1.82 | 1.25 | 0.491 | 2.40 | 1.77 |
| *df* | 1, 43.6 | 1, 43.4 | 1, 43.5 | 1, 35.5 | 1, 35.3 | 1, 35.5 | 1, 35.3 |
| **W × S** |  |  |  |  |  |  |  |
| *P*-value | **0.530** | **0.687** | **0.774** | **0.357** | **0.298** | **0.741** | **0.144** |
| *F*-value | 0.401 | 0.164 | 0.084 | 0.871 | 1.11 | 0.111 | 2.23 |
| *df* | 1, 44.0 | 1, 43.8 | 1, 43.9 | 1, 35.5 | 1, 35.2 | 1, 35.5 | 1, 35.3 |
| **I × W × S** |  |  |  |  |  |  |  |
| *P*-value | **0.126** | **0.198** | **0.154** | **0.853** | **0.497** | **0.321** | **0.993** |
| *F*-value | 2.44 | 1.71 | 2.10 | 0.035 | 0.471 | 1.01 | 0.0001 |
| *df* | 1, 43.7 | 1, 43.6 | 1, 43.6 | 1, 35.5 | 1, 35.4 | 1, 35.5 | 1, 35.4 |

**P**, **F** and **df** values are in **bold**, *italics* and regular type, respectively, significance codes: <0.0001-0.001'\*\*\*\*', 0.001-0.01'\*\*\*', 0.01-0.05'\*\*. .

**FIGURE 4.**
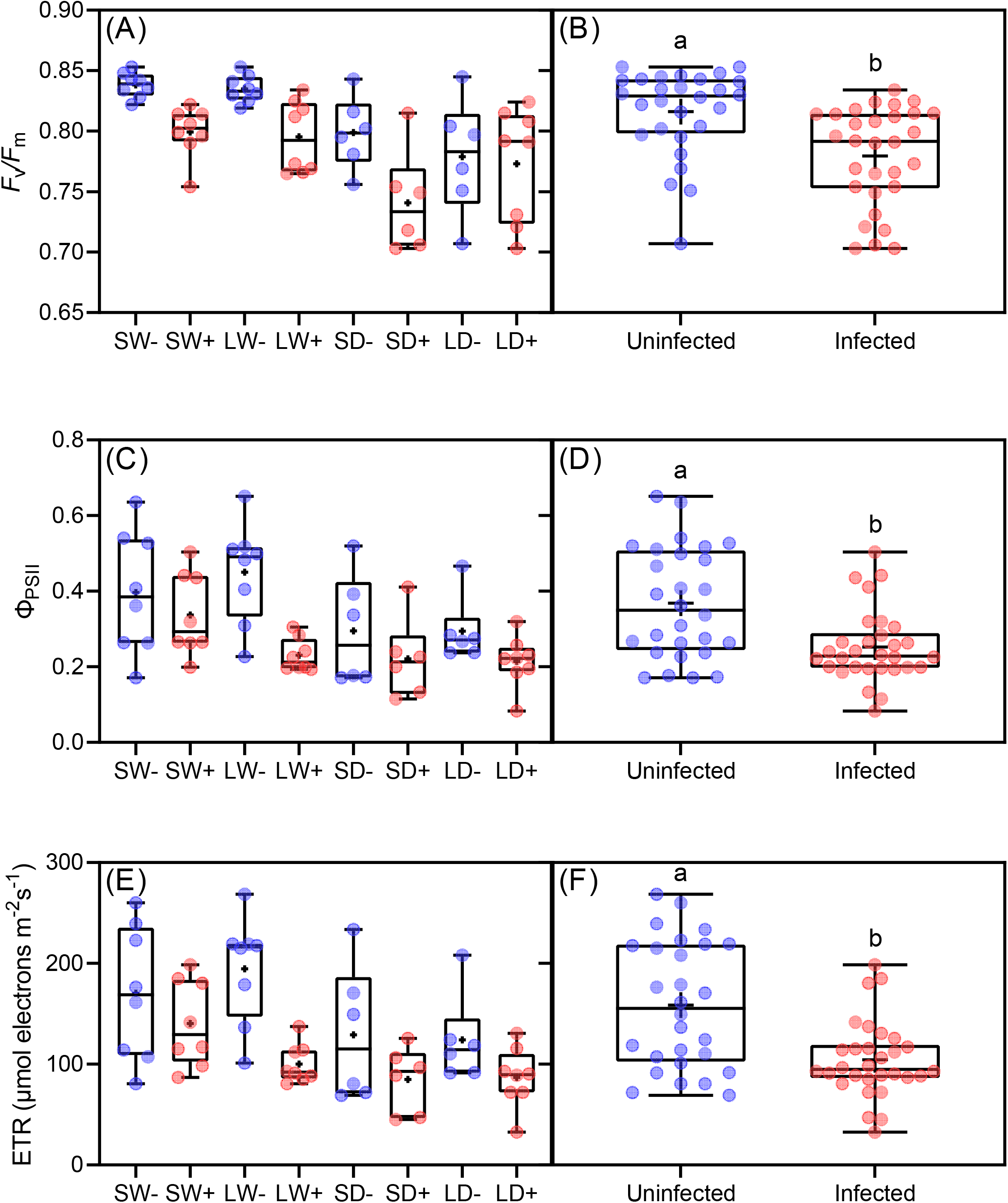
(A) Predawn and (C) midday quantum yield (*F*_v_/*F*_m_, Φ_PSII_) and (E) midday electron transport rates (ETR) of small (S) or large (L) *Ulex europaeus* uninfected (-) or infected (+) with *Cassytha pubescens* growing in well-watered (W) or low water (D) conditions. Main effect of infection on parasite (B) *F*_v_/*F*_m_, (d) Φ_PSII_ and (F) ETR. All data points, median, 1^st^ and 3^rd^ quartiles, interquartile range and mean (+ within box) are shown, different letters signify significant differences: (A, C, E) *n* = 8 (except SD−, LD− and SD+: *n* = 6) and (B, D, F) *n* = 28–30

**FIGURE 5.**
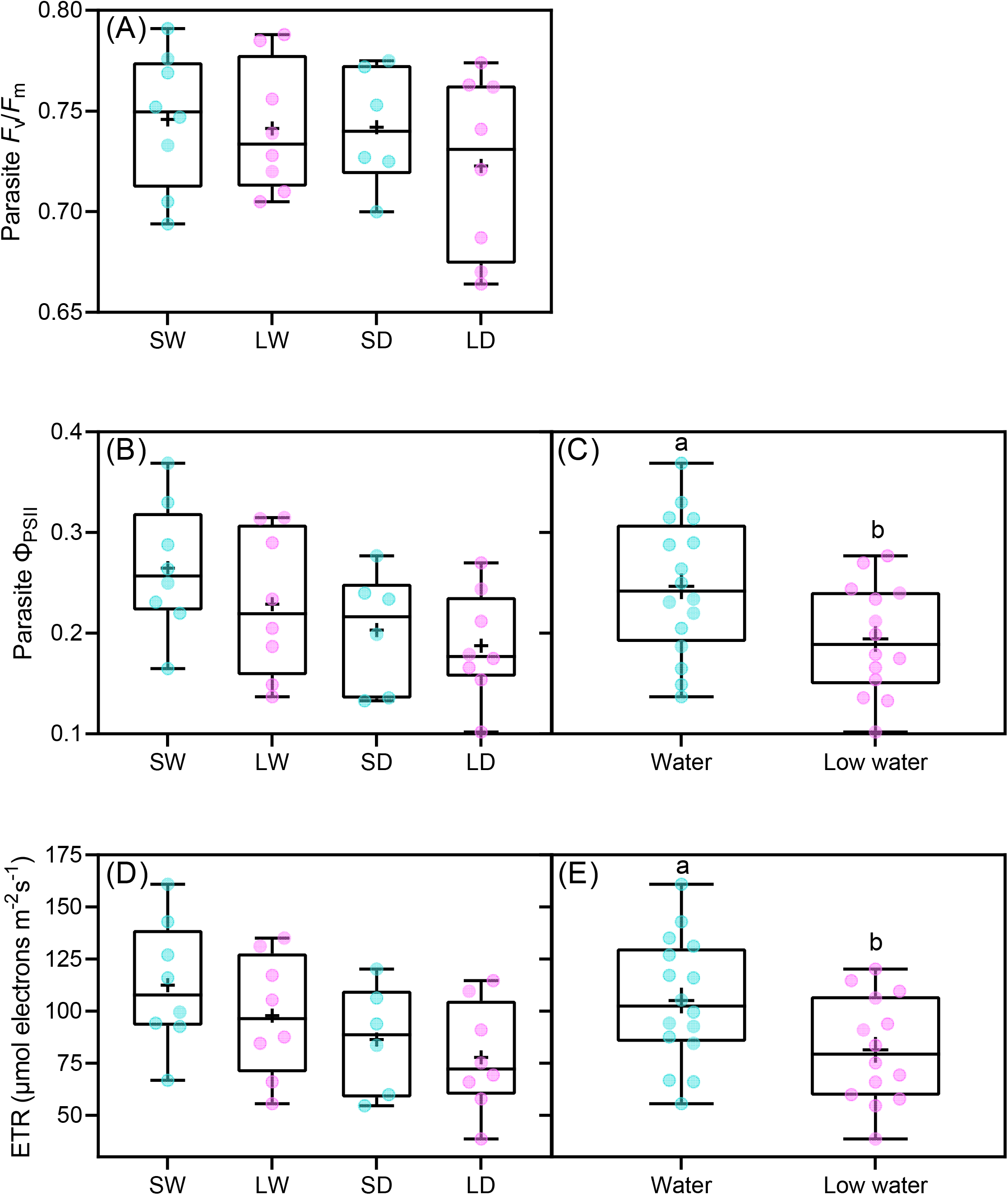
(A) Predawn and (B) midday quantum yield (*F*_v_/*F*_m_, Φ_PSII_) and (D) midday electron transport rates (ETR) of *Cassytha pubescens* when infecting small (S) or large (L) *Ulex europaeus* growing in well-watered (W) or low water (D) conditions. Main effect of water on parasite (C) Φ_PSII_ and (E) ETR. All data points, median, 1^st^ and 3^rd^ quartiles, interquartile range and mean (+ within box) are shown, different letters signify significant differences: (A, B, D) *n* = 8 (except SD: *n* = 6) and (C, E) *n* = 14–16

There was a main effect of water and host size on δ^13^C of *U. europaeus* (Table 3, no threeway interaction: Figure 6A). δ^13^C of low water plants was significantly higher than that of well-watered plants (Figure 6B). δ^13^C of large plants was significantly higher than that of small plants (Figure 6C). When comparing between δ^13^C of infected plants and associated parasite, we detected interactive effects of species × water (*F*_1, 34.2_ = 11.9; *P* = 0.002) and species × host size (*F*_1, 34.2_ = 5.25; *P* = 0.028) (no three-way interaction: Figure 6D). In well-watered and low water conditions, δ^13^C of *C. pubescens* was 1.8‰ and 2.7‰ higher than that of respective hosts (Figure 6E). For small and large hosts, δ^13^C of *C. pubescens* was 2.6‰ and 2.0‰ higher than that of respective hosts (Figure 6F). There was a main effect of water on δ^13^C of *C. pubescens*, which was 1.3‰ higher in low water than in well-watered conditions (Table 2; Figure 6E).

**FIGURE 6.**
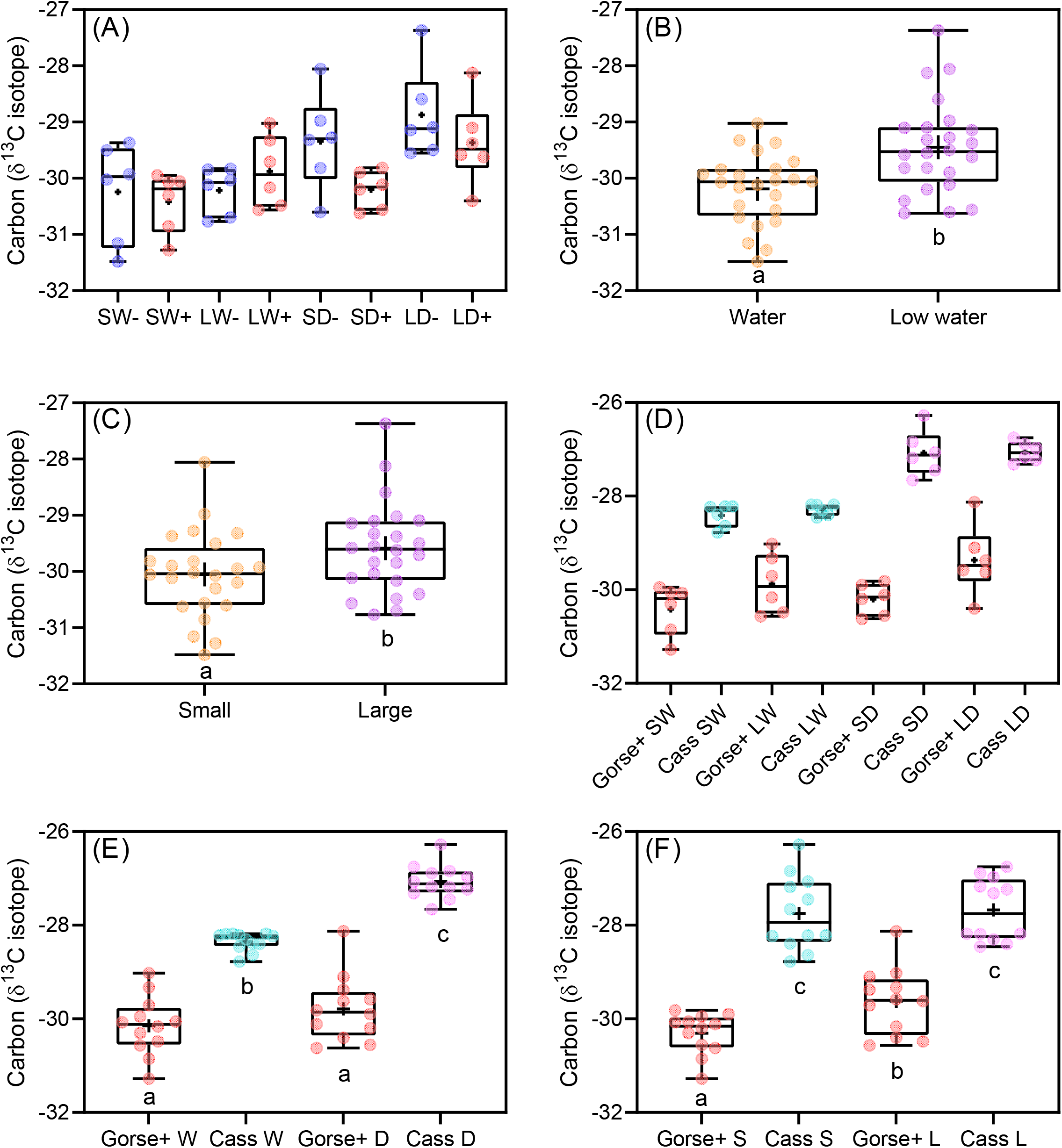
(A) Carbon isotope composition (δ^13^C ‰) of small (S) or large (L) *Ulex europaeus* uninfected (-) or infected (+) with *Cassytha pubescens* growing in well-watered (W) or low water (D) conditions. Main effect of (B) water and (C) size on host δ^13^C. (D) δ^13^C of infected host and associated parasite. (E) Species × water and (F) species × size interaction on comparison between δ^13^C of infected host (Gorse +) and associated parasite (Cass). All data points, median, 1^st^ and 3^rd^ quartiles, interquartile range and mean (+ within box) are shown, different letters signify significant differences: (A, D) *n* = 6, (B, C) *n* = 24 and (E, F) *n* = 12

Infection and water had an interactive effect on foliar [N] and [K] of *U. europaeus* (Table 3, no three-way interactions: Figure 7A,C). Foliar [N] and [K] of infected plants in well-watered conditions were 11% and 29% lower than those of respective uninfected plants while [N] and [K] of low water plants were unaffected by infection (Figure 7B,D). There was a main effect of infection and host size on [P] of *U. europaeus* (Table 3, no three-way interaction: Figure 7E). Foliar [P] of infected plants was 23% lower compared with that of uninfected plants (Figure 7F). Foliar [P] of large plants plants was 24% lower relative to that of small plants (Figure S8A). No significant effects were found for parasite [N] (Table 2; Figure 8A). There was a main effect of water on parasite [K] and [P] (Table 2, no two-way interactions: Figure 8B,D). Stem [K] and [P] of *C. pubescens* in well-watered conditions were 23% and 15% lower than those in low water treatments (Figure 8C,E). There was also a main effect of host size on parasite [P] which for *C. pubescens* when infecting large hosts was 15% lower relative to that when infecting small hosts (Table 2; Figure S8B).

**FIGURE 7.**
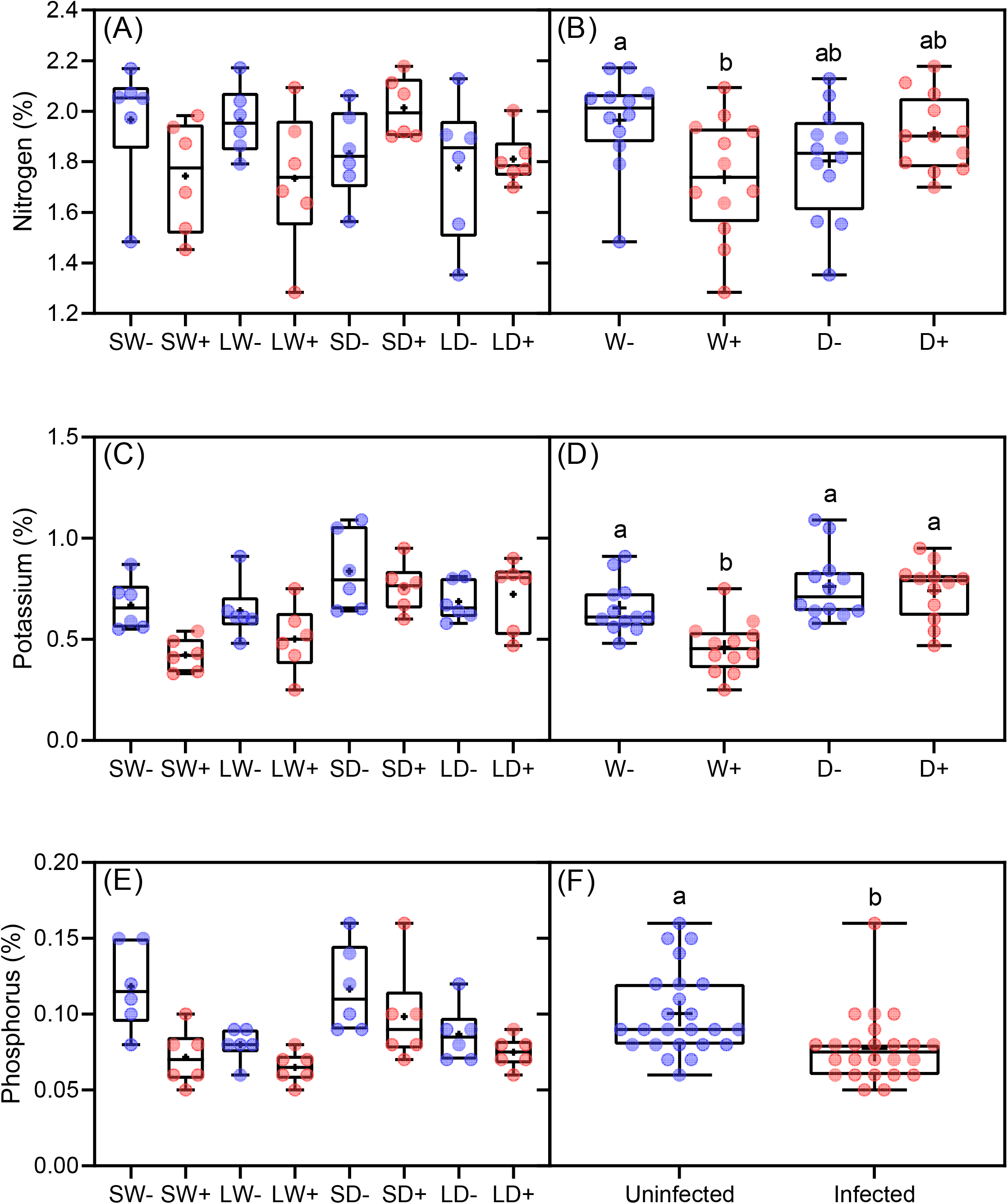
(A) Foliar nitrogen, (C) potassium and (E) phosphorus concentration of small (S) or large (L) *Ulex europaeus* uninfected (-) or infected (+) with *Cassytha pubescens* growing in well-watered (W) or low water (D) conditions. Infection × water interaction on host (B) nitrogen and (D) potassium. (F) Main effect of infection on host phosphorus. All data points, median, 1^st^ and 3^rd^ quartiles, interquartile range and mean (+ within box) are shown, different letters signify significant differences: (A, C, E) *n* = 6, (B, D) *n* = 12 and (F) *n* = 24

**FIGURE 8.**
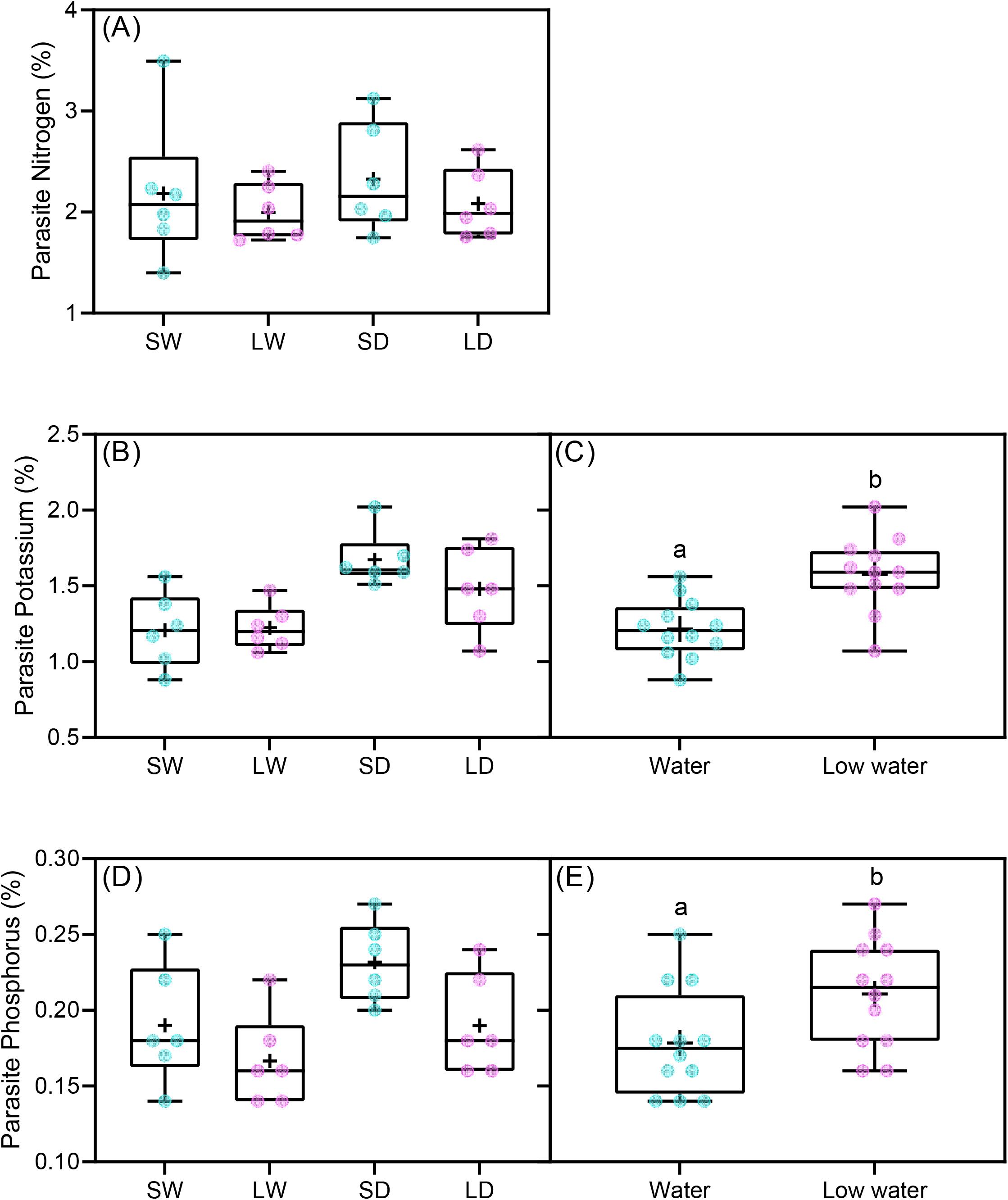
(A) Stem nitrogen, (B) potassium and (D) phosphorus concentration of *Cassytha pubescens* when infecting (S) or (L) *U. europaeus* growing in well-watered (W) or low water (D) conditions. Main effect of water on parasite (C) potassium and (E) phosphorus concentrations. All data points, median, 1^st^ and 3^rd^ quartiles, interquartile range and mean (+ within box) are shown, different letters signify significant differences: (A, B, D) *n* = 6, and (C, E) *n* = 12

## Discussion

Our hypotheses that small well-watered plants would suffer most from infection and the parasite would grow significantly more on large well-watered plants were not supported by the data. However, we did find that the strong negative effect of *C. pubescens* on total biomass of *U. europaeus* was more severe in: 1) well-watered conditions and 2) when the host was smaller. We also found that the parasite had more biomass in well-watered conditions or when the host was larger. In addition, infection or low water conditions strongly suppressed host photosynthetic performance.

Importantly, we found that infection had a more severe negative effect on biomass of *U. europaeus* in well-watered than in low water conditions as similarly found in Cirocco et al. (2016b, 2021b). The holoparasitic vine *Cuscuta gronovii* also had a stronger negative effect on the growth of *Verbesina alternifolia* in well-watered than in drought conditions (Evans and Borowicz, 2013). In contrast, the root hemiparasite *Rhinanthus alectorolophus* had a strong negative impact on host growth, regardless of water treatments, including under varying nitrogen conditions (Světlíková et al., 2018; Korell et al., 2020). On the other hand, Těšitel et al. (2015) found that *R. alectorolophus* had the greatest negative impact on host biomass in low water and high nitrogen conditions, possibly due to the parasite being able to maintain effective resource removal (via high transpiration rates) despite incurring low water conditions. *Cassytha pubescens* does not seem to share this ability as inferred from: 1) its much higher δ^13^C than the host in low water conditions (Figure 6E) and 2) that it visibly wilted when the host incurred field capacities below 60% which is likely due to it being a thin vine that is prone to desiccation (Cirocco pers. obs.). Here, the greater parasite vigour on well-watered hosts, likely explains why the parasite was able to grow better (when standardised for host growth, Figure 3E) and remove more resources and have a greater impact on host growth in these conditions. This is corroborated by the fact that infection strongly decreased host foliar [N] and [K] only in the well-watered treatment (Figure 7B,D).

Other studies have also found improved parasite growth in well-watered conditions for associations involving *C. pubescens-U. europaeus, Cuscuta gronovii-Verbesina alternifolia* and *R. alectorolophus* when infecting a variety of hosts (Evans and Borowicz, 2013; Cirocco et al., 2016b; Světlíková et al., 2018; Korell et al., 2020). In contrast, Cirocco et al. (2021b) found that the *C. pubescens* grew similarly on *U. europaeus* in both high and low water conditions, irrespective of nitrogen treatments. Water deficits imposed in Cirocco et al. (2021b) may not have been low enough to elicit similar responses to those found here for parasite growth. A study by Těšitel et al. (2015) found that the growth of *R. alectorolophus* improved in high water conditions (when N supply was low), however, the parasite achieved the greatest biomass (strongly correlated with parasite *F*_v_/*F*_m_) in high N conditions when water supply was either high or low. Here, improved parasite growth in well-watered conditions may also be partly due to enhanced photosynthetic performance (Φ_PSII_ and ETR, but not *F*_v_/*F*_m_) of *C. pubescens* when growing on well-watered hosts. Similarly, Cirocco et al. (2016b) found that *F*_v_/*F*_m_ (but not Φ_PSII_) of *C. pubescens* was significantly higher in well-watered compared with low water treatments. In contrast, Cirocco et al. (2021b) found that photosynthetic performance (*F*_v_/*F*_m_, Φ_PSII_ or ETR) of *C. pubescens* was unaffected by water or nitrogen supply. As mentioned, this difference between findings may be due to the low water treatment imposed in Cirocco et al. (2021b) not being low enough (same field capacity: 60% as our study, but much larger pots) to elicit a similar response (i.e. suppressed parasite photosynthetic performance).

Another key finding was that infection had a stronger negative effect on total biomass of smaller than larger *U. europaeus*, as similarly found in Cirocco et al. (2020). Li et al. (2015) also found that *Cuscuta australis* negatively affected the total biomass of younger, smaller *Bidens pilosa*, whereas older, larger hosts were unaffected by the parasite. As we found larger plants to also be affected by infection, we assume this difference between studies may be due to *C. pubescens* negatively affecting host photosynthetic performance regardless of host size, whereas *Cuscuta australis* only negatively affected photosynthesis of younger hosts. In our study, as the parasite more strongly affected growth of small hosts relative to large hosts by only 4%, it is understandable that there was no data herein that could be help explain this infection × size interaction. Nevertheless, we can speculate that smaller plants have less resources to buffer the effects of resource removal from the parasite and that any parasite-induced decreases in nutrients of *U. europaeus* may have been masked by concurrent decreases in host growth (Cirocco et al., 2021a).

Along with removal of resources, the parasite had a significant negative effect on host *F*_v_/*F*_m_, regardless of water treatments or host size as similarly found in Cirocco et al. (2016b, 2020). Cirocco et al. (2021b) also found that *C. pubescens* negatively affected *F*_v_/*F*_m_ of *U. europaeus*, irrespective of water or nitrogen supply. In contrast, Těšitel et al. (2015) found that *R. alectorolophus* negatively affected *F*_v_/*F*_m_ of maize (but not wheat) only when water was limiting and nitrogen supply was abundant. This disparity between findings again might in part be related to *R. alectorolophus* (but not *C. pubescens*) being able to maintain high stomatal conductance and transpiration rates when incurring low water conditions. This action would maintain a water potential gradient that favours the movement of resources from the host to the parasite (Těšitel et al., 2015). In other studies, *C. pubescens* has also been found to negatively affect *F*_v_/*F*_m_ of the invasive host *Cytisus scoparius* but generally not that of the native hosts, *Leptospermum myrsinoides* and *Acacia paradoxa* (Shen et al., 2010; Cirocco et al., 2015, 2021a; but see Prider et al., 2009 and the latter study). Here, the negative effect of the parasite on *F*_v_/*F*_m_ of *U. europaeus* may be due to host ETR being adversely affected by infection (Figure 4F) likely via resource removal, namely phosphorus (Figure 7F) (Rychter and Rao, 2005). Electron transport rates can be used as a proxy for photosynthesis and its decline would increase the ratio of PFD to photosynthesis thereby exposing the host to potentially prolonged periods of excess light, and consequent chronic photoinhibition as reflected by suppressed *F*_v_/*F*_m_ (Demmig-Adams and Adams, 2006).

## Conclusion

Our results support the potential-use of some native parasites as biological agents for weeds, and suggest that they would be especially effective in controlling invasive hosts and conserving biodiversity in wetter habitats that have been recently invaded (i.e. hosts smaller in size). Our results also suggest that some parasites will still have a strong negative effect on growth of invasive hosts in low water conditions, which are expected to become more frequent, especially in Mediterranean systems as a result of climate change. Noteworthy is the similarity between parasites of similar growth form (hemi- and holoparasitic vines) in terms of improved growth in well-watered conditions and greater impact when hosts are well-watered or smaller. More studies are needed, however, on root hemiparasites before more meaningful comparisons can be made, considering that there are no studies which have investigated root parasites in association with hosts of different size or age. Moreover, our findings set a global baseline for comparison with other studies investigating how the combined effects of water supply and host size influence the parasite:host association.

## Author Contributions

RMC and JMF conceived and designed the experiment. RMC performed the experiment and analysed the data. RMC, JMF, and EF interpreted the analysis and wrote the manuscript.

## Acknowledgements

Special thanks to Dr Tony Hall (Mawson Analytical Spectrometry Services, The University of Adelaide) for his expert analysis with the IRMS. This work was supported by the School of Biological Sciences Early and Mid-Career Researcher Association Small Grant Scheme (The University of Adelaide) [15125342].

## Supporting information

**Fig. S1.** Height of plants before and after infection process.

**Fig. S2-S5.** Photos of plants at the end of the experiment.

**Fig. S6.** Water × size interaction and size effect on host growth measures.

**Fig. S7.** Water effect on host photosynthetic performance.

**Fig. S8.** Host size effect on host and parasite phosphorus concentration.

