## Supplementary material for "Do water and host size interactively affect the impact of a native hemiparasite on a major invasive legume?": Supp Figs S1-S8

**Fig. S1.** Height of plants before and after infection process.

**Fig. S2-S5.** Photos of plants at the end of the experiment.

**Fig. S6.** Water × size interaction and size effect on host growth measures.

**Fig. S7.** Water effect on host photosynthetic performance.

**Fig. S8.** Host size effect on host and parasite phosphorus concentration.

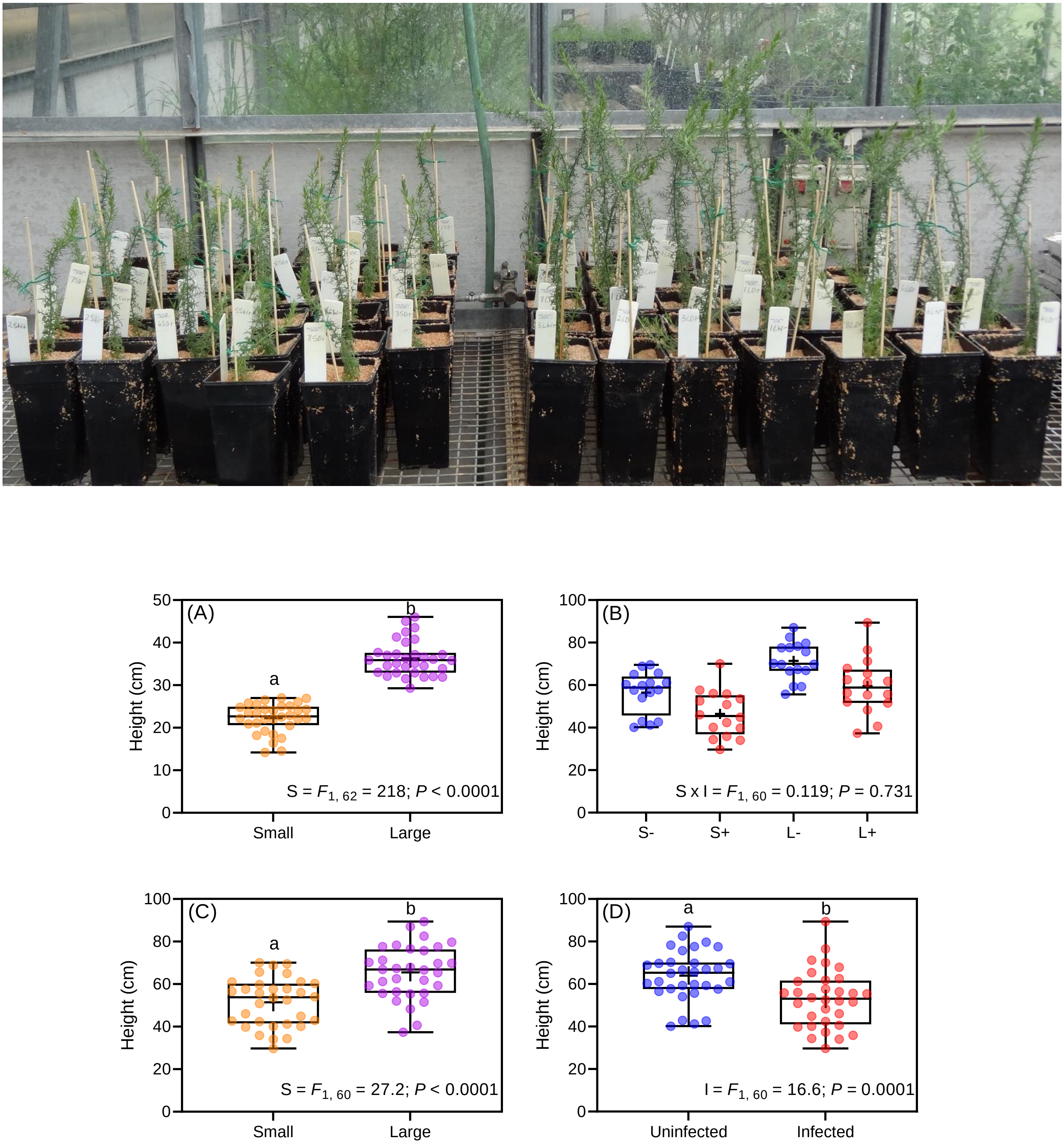


**FIGURE S1** Photo of small and large *U. europaeus* before the infection process in late October

2020. Height of small (S) and large (L) *Ulex europaeus* before (A) and after (B) the infection process when uninfected (–) and infected plants (+) were obtained. Main effects of (C) size and (D) infection with *Cassytha pubescens* on height of *U. europaeus* once the infection process was completed. All data points, median, 1st and 3rd quartiles, interquartile range and mean (+ within box) are displayed and different letters denote significant differences: (A, C, D) $n$ = 32 and (b) $n$ = 9. Statistical output for the effect of size (S), infection (I) and their interaction (S × I) on plant height are shown in figure panels

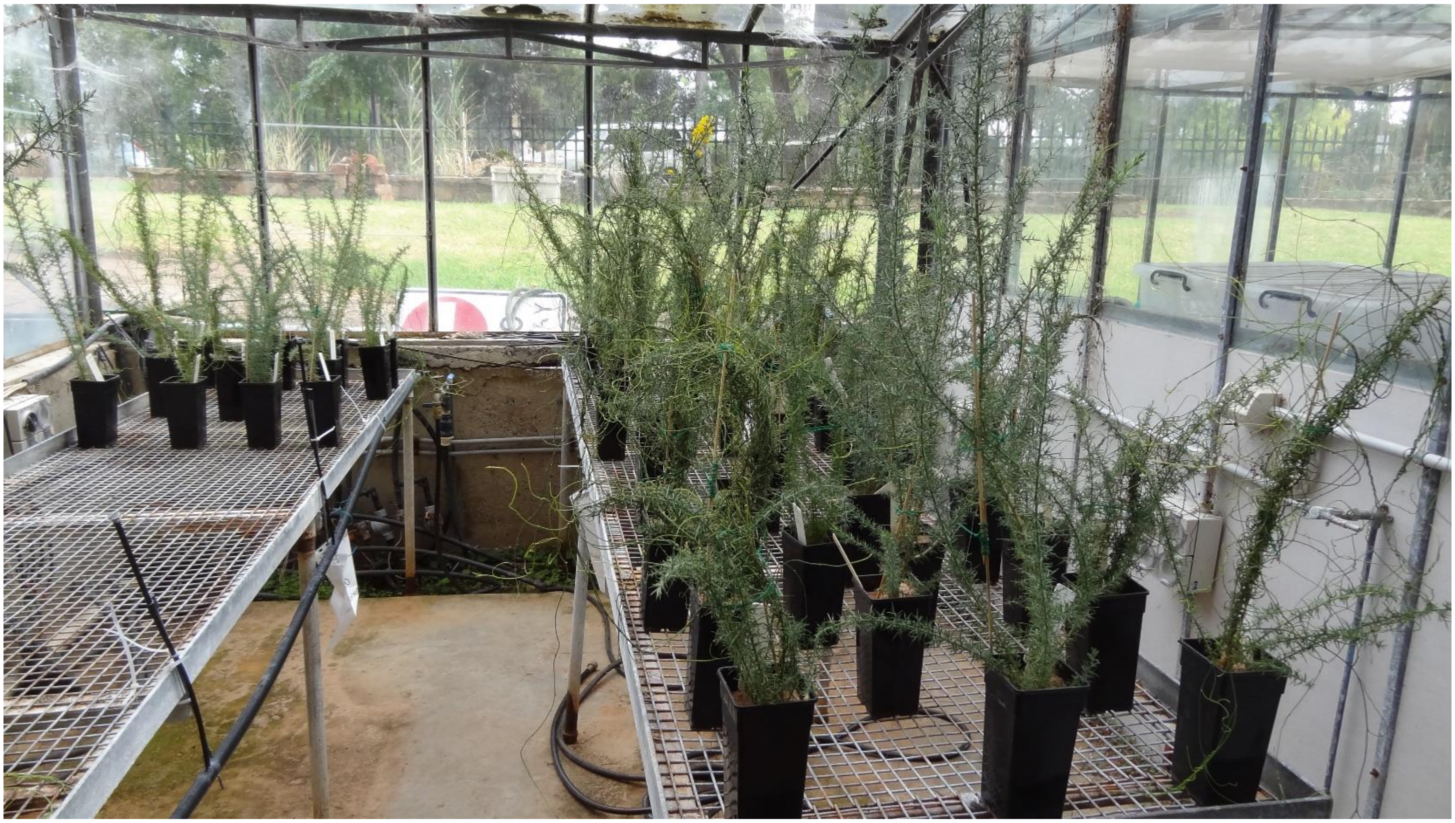

**FIGURE S2** Photo of experimental plants at the end of the experiment in early April 2021

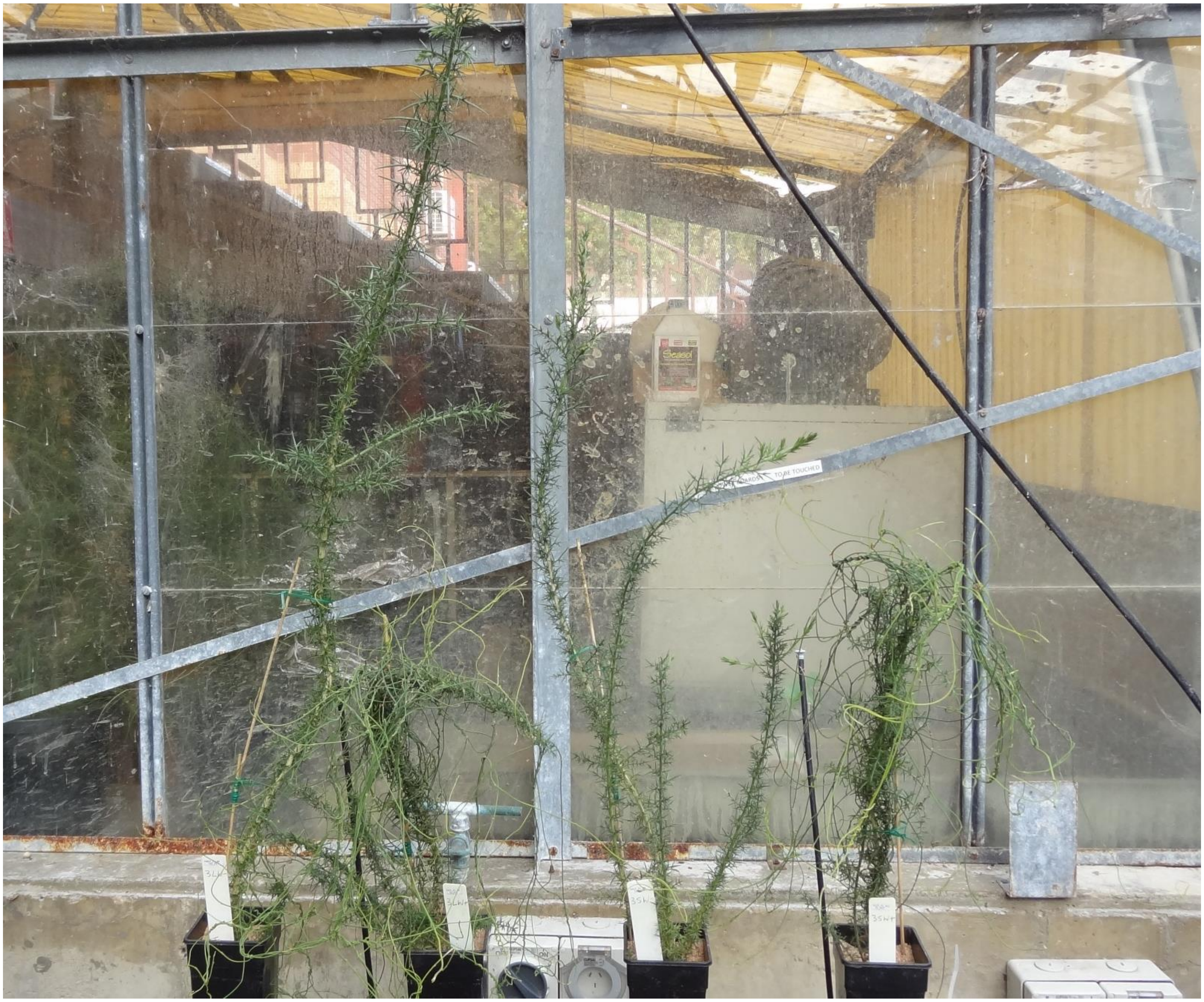

**FIGURE S3** Photo taken at the end of the experiment in early April 2021 of large (L) and small (S) uninfected (–) and infected (+) *U. europaeus*, well-watered (W) from Block 3, left to right: LW–, LW+, SW– and SW+

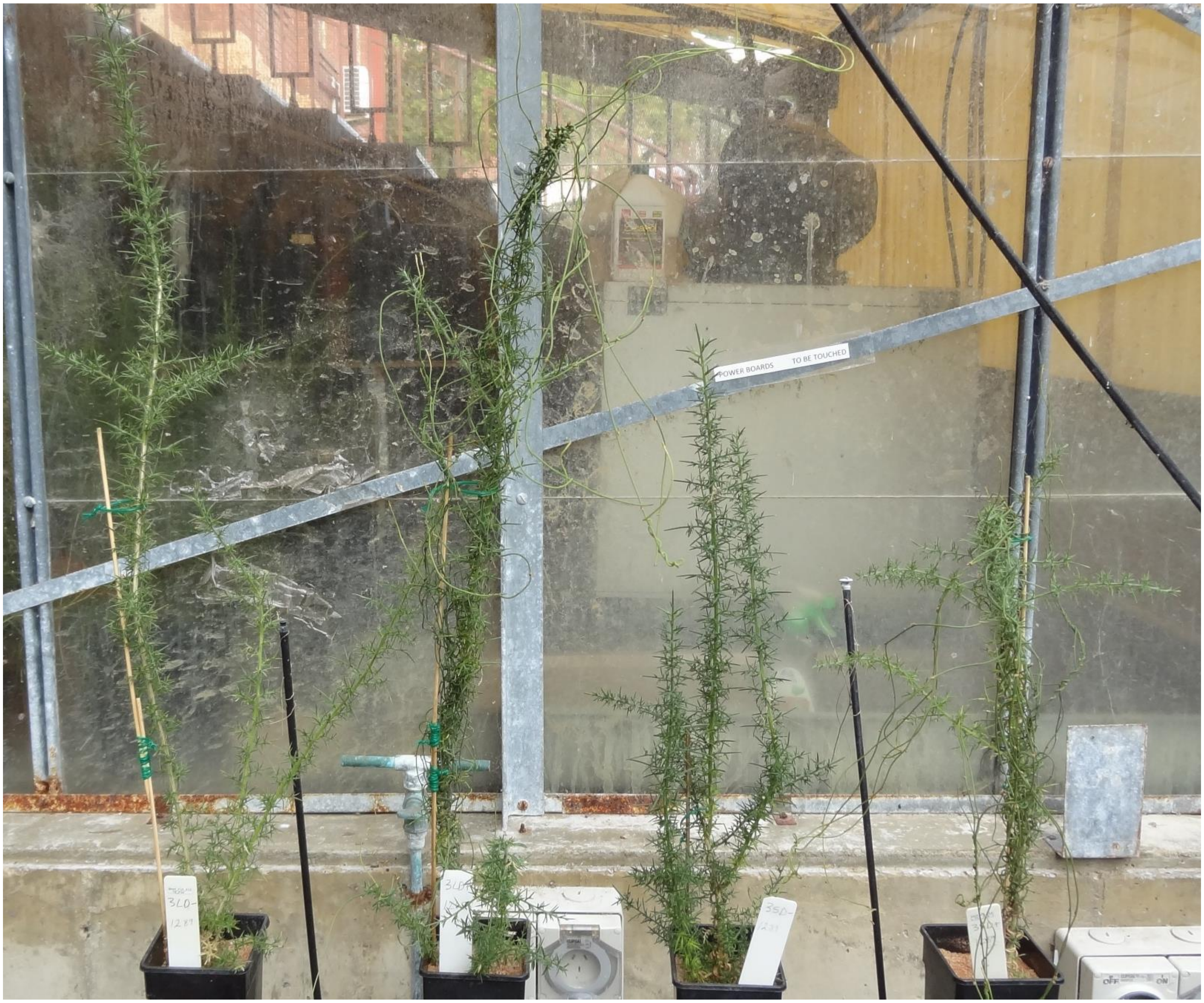


**FIGURE S4** Photo taken at the end of the experiment in early April 2021 of large (L) and small (S) uninfected (–) and infected (+) *U. europaeus* supplied with low water (D) from Block 3, left to right: LD–, LD+, SD– and SD+

(A)

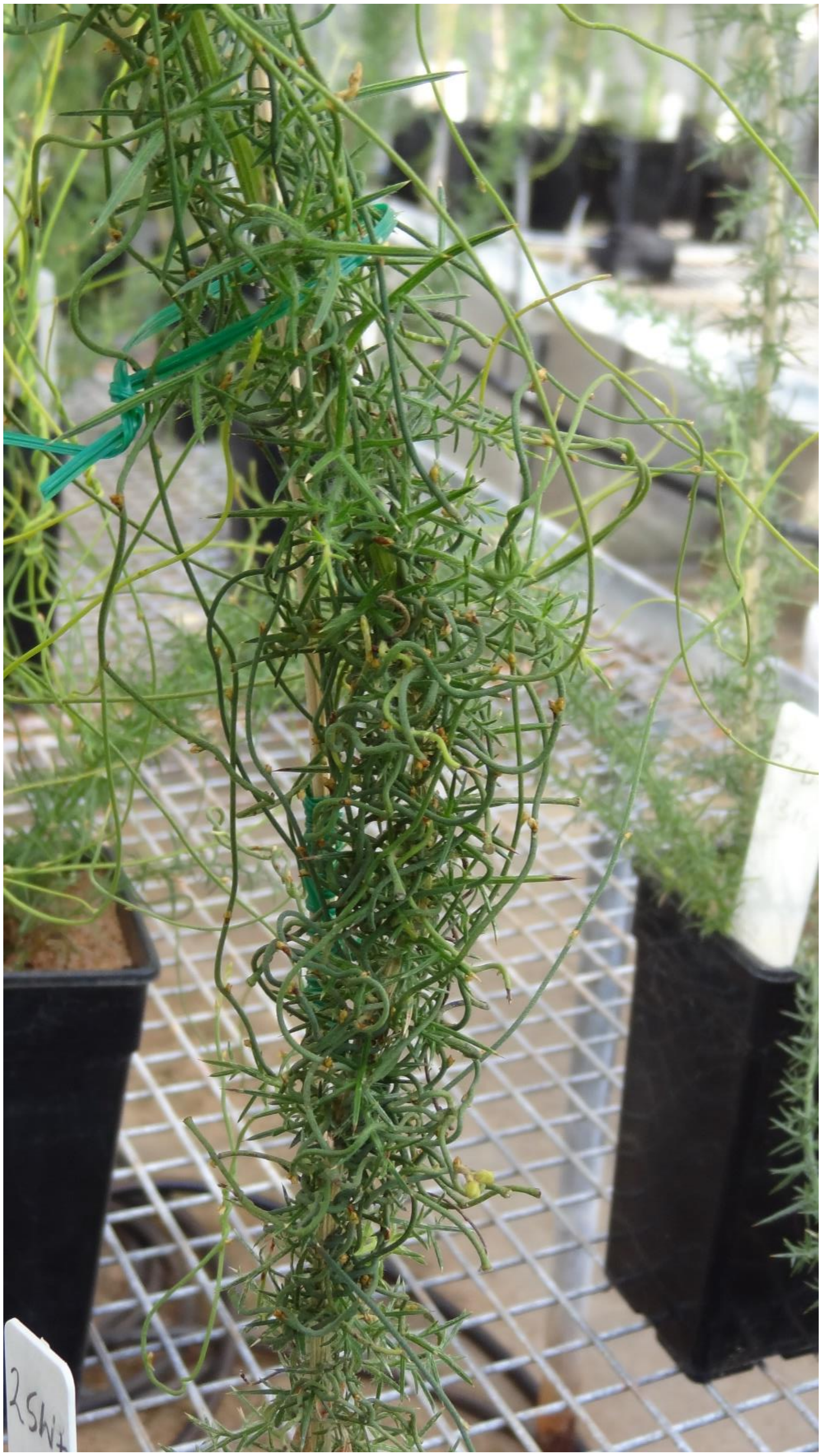

(B)

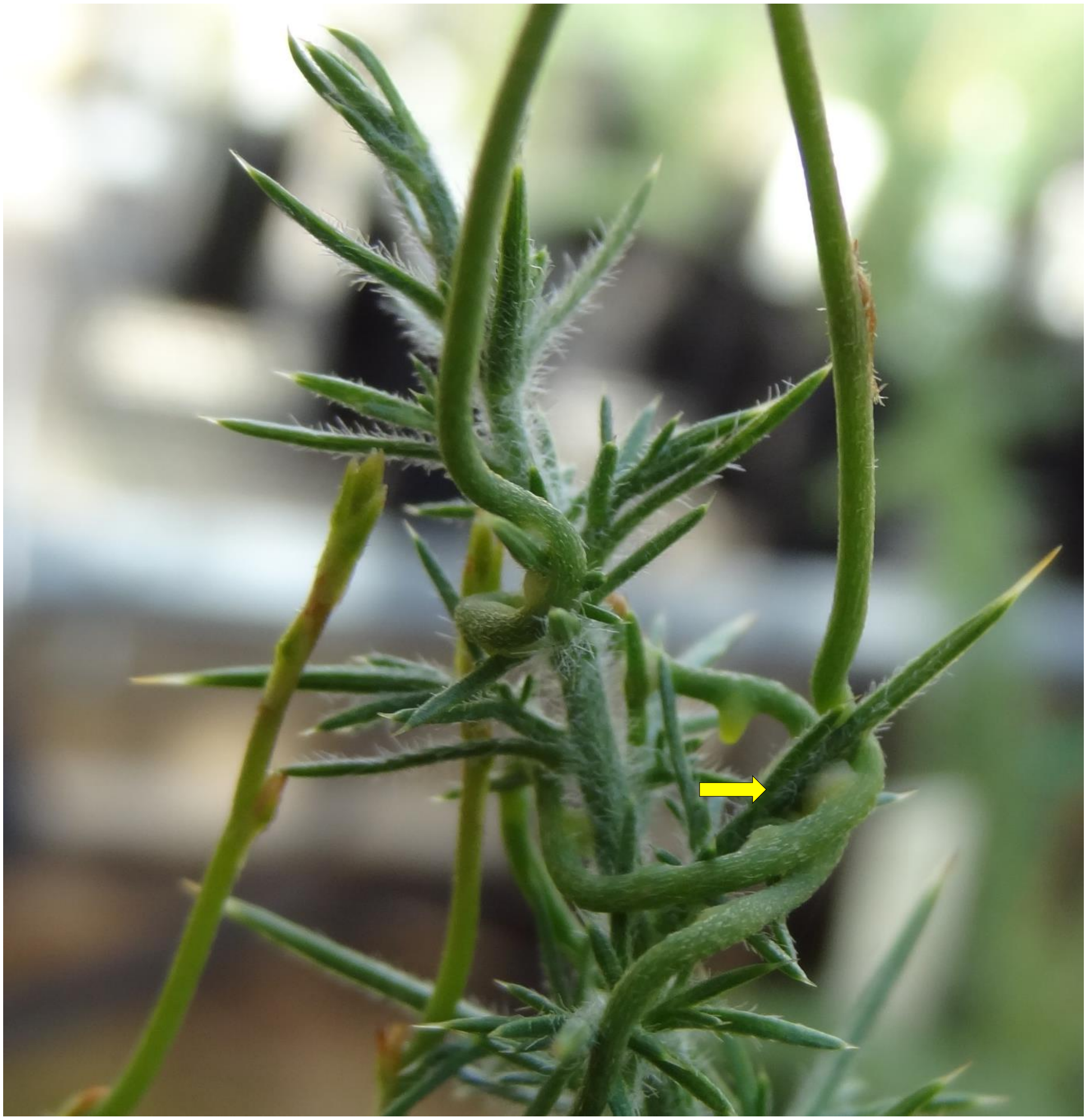

**FIGURE S5** Photo taken at the end of the experiment in early April 2021 of (A) small, well-watered *Ulex europaeus* infected with *Cassytha pubescens* and (B) close-up of the parasite's haustoria attached to a host spine (yellow arrow)

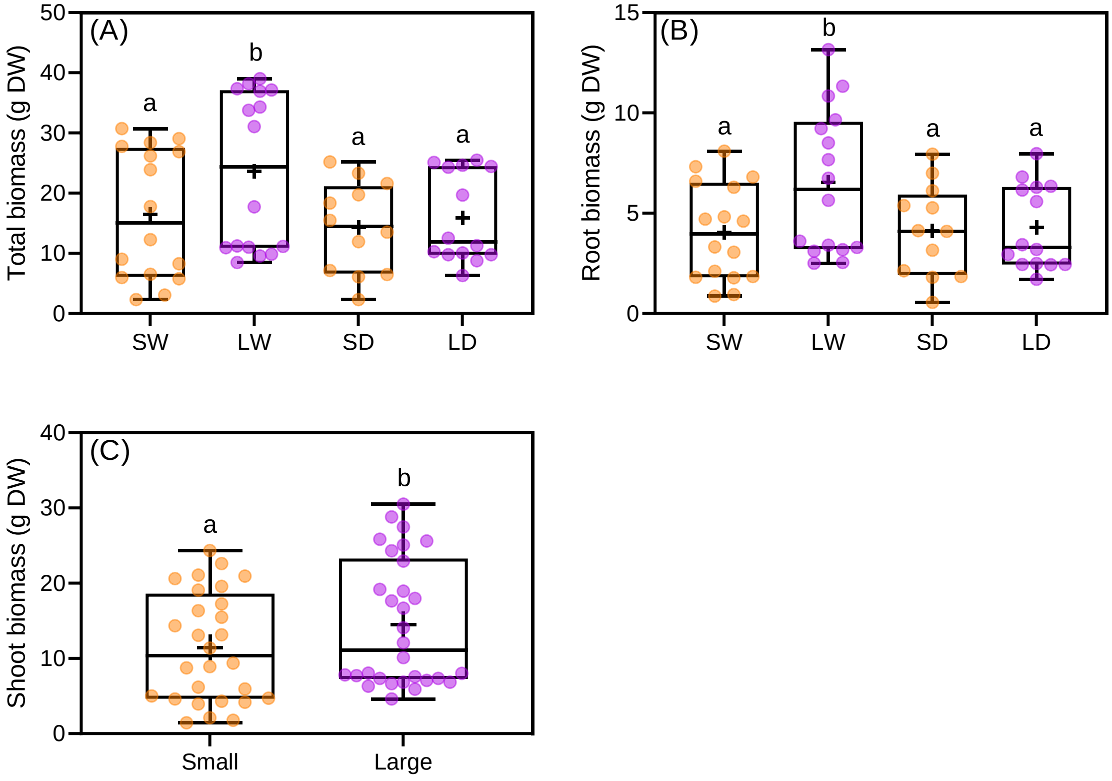


**FIGURE S6** Water × size interaction for (A) total, and (B) root biomass of small (S) or large (L) *Ulex europaeus* growing in well-watered or low water conditions (W=uninfected and infected well-watered plants pooled; D=uninfected and infected low-watered plants pooled). (C) Main effect of host size on shoot biomass of *U. europaeus*. All data points, median, 1st and 3rd quartiles, interquartile range and mean (+ within box) are shown, different letters signify significant differences: (A, B) *n* = 16 (except SD and LD: *n* = 12 and 14, respectively) and (C) *n* = 28–30

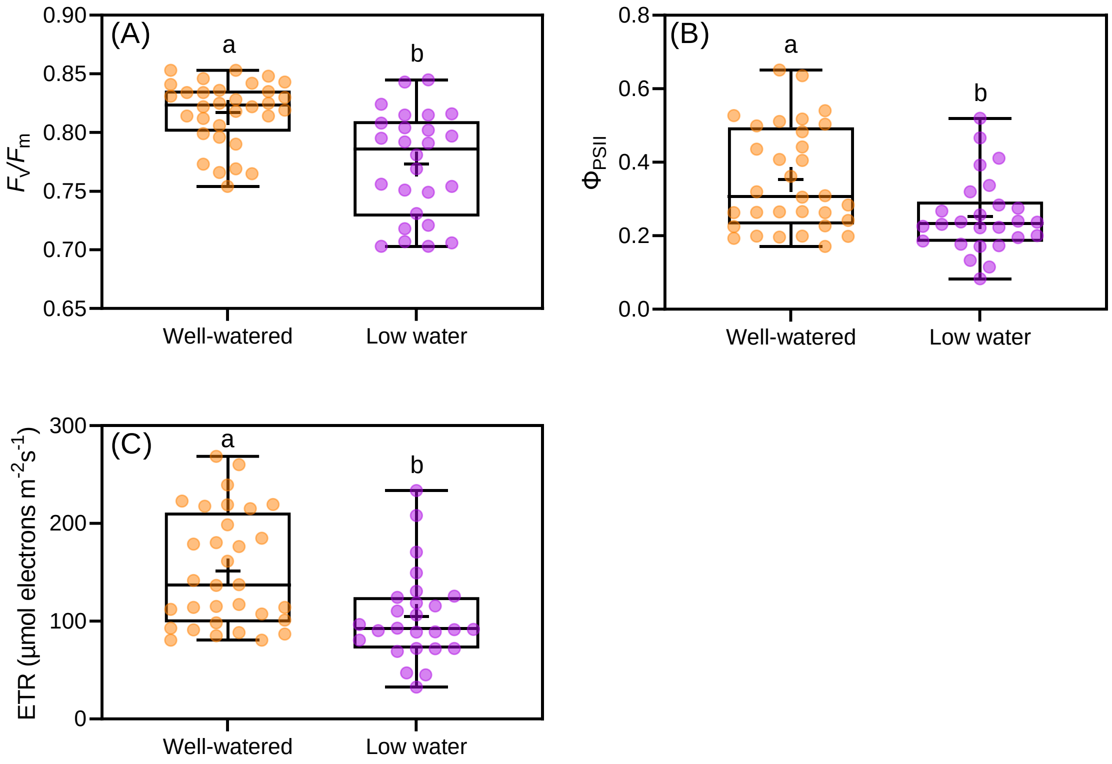


**FIGURE S7** Main effect of water on (A) predawn and (B) midday quantum yield ($F_v/F_m$, $\Phi_{PSII}$) and (C) midday electron transport rates (ETR) of *Ulex europaeus* (uninfected and infected small and large well-watered plants pooled compared with uninfected and infected small and large low water plants pooled). All data points, median, 1$^{st}$ and 3$^{rd}$ quartiles, interquartile range and mean (+ within box) are shown, different letters signify significant differences: (A, B, C) $n$ = 32 (well-watered) and $n$ = 26 (low water)

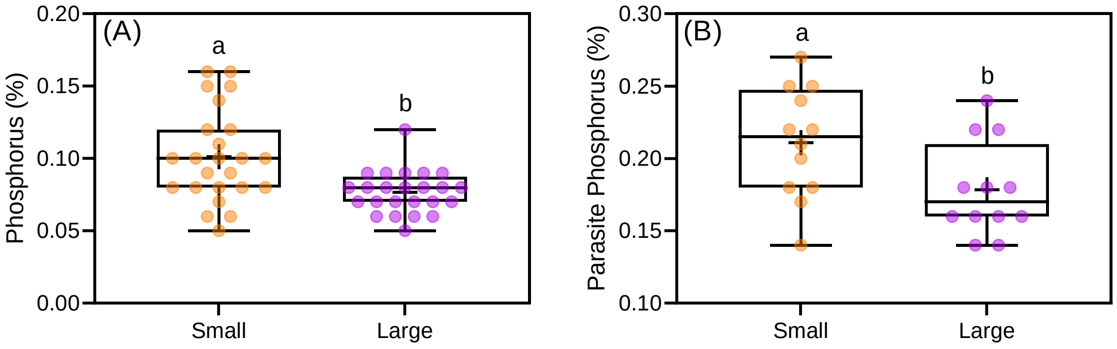


**FIGURE S8** (A) Main effect of host size on phosphorus concentration of *Ulex europaeus* (small uninfected and infected plants in well-watered and low water treatments pooled compared with large uninfected and infected plants in well-watered and low water treatments pooled). (B) Main effect of host size on phosphorus concentration of *Cassytha pubescens* (parasite growing on small plants in well-watered and low water treatments pooled compared with parasite growing on large plants in well-watered and low water treatments pooled). All data points, median, 1$^{st}$ and 3$^{rd}$ quartiles, interquartile range and mean (+ within box) are shown, different letters signify significant differences: (A) $n$ = 24 and (B) $n$ = 12
